# Hierarchical and scaffolded phosphorylation of two degrons controls PER2 stability

**DOI:** 10.1101/2023.11.02.565258

**Authors:** Joel C. Francisco, David M Virshup

## Abstract

The duration of the transcription-repression cycles that give rise to mammalian circadian rhythms is largely determined by the stability of the PERIOD protein, the rate-limiting components of the molecular clock. The degradation of PERs is tightly regulated by multisite phosphorylation by Casein Kinase 1 (CK1δ/ε). In this phosphoswitch, phosphorylation of a PER2 degron (Degron 2, D2) causes degradation, while phosphorylation of the PER2 Familial Advanced Sleep Phase (FASP) domain blocks CK1 activity on the degron, stabilizing PER2. However, this model and many other studies of PER2 degradation do not include the second degron of PER2 that is conserved in PER1, termed Degron 1, D1. We examined how these two degrons contribute to PER2 stability, affect the balance of the phosphoswitch, and how they are differentiated by CK1. Using PER2-luciferase fusions and real-time luminometry, we investigated the contribution of both D2 and of CK1-PER2 binding. We find that D1, like D2, is a substrate of CK1 but that D1 plays only a ‘backup’ role in PER2 degradation. Notably, CK1 bound to a PER1:PER2 dimer protein can phosphorylate PER1 D1 in trans. This scaffolded phosphorylation provides additional levels of control to PER stability and circadian rhythms.

## Introduction

Daily rhythmic oscillations in biological processes are regulated by the circadian clock, a molecular time-keeper intrinsic to most eukaryotes. The duration of the transcription-repression cycles that give rise to mammalian circadian rhythms are determined in large part by the stability of the PERIOD proteins PER1 and PER2, rate-limiting components of the molecular clock (1–5). As such, the degradation of PERs is a tightly regulated process. Circadian period-altering mutations in Drosophila (6), in hamsters (7, 8), and in humans (9) identified Casein Kinase 1ε (CK1ε) and its paralog CK1δ as critical determinants of PER stability (8, 10–12). Mutations in diverse organisms that result in altered PER phosphorylation by CK1 result in profound dysregulation of the circadian clock (6, 9, 10, 13, 14).

The importance of the CK1 phosphorylation of PER in maintaining robust circadian rhythms is also reflected in the conserved complexity of their interaction and regulation (15, 16). CK1 interacts with PER proteins via two CK1 Binding Domains (CKBD) (1, 11). Disruption of either CKBD-A or -B compromised CK1 binding with PER2 thereby altering PER2 stability (11, 17). In mammals, the first critical phosphorylation site on PER2 was identified in a family with Familial Advanced Sleep Phase (FASP)(13, 18). FASP is characterized by early sleep onset and a short circadian period. Affected individuals have a S662G mutation in PER2 (10). Normally, phosphorylation of PER2 S662 (S659 in mouse) primes the downstream serine residues for rapid phosphorylation (10, 19, 20). The S662G mutation thus also prevents phosphorylation of PER2 at serines 665, -668, -671, and -674 and significantly destabilizes the protein, causing the short circadian period phenotype (10). This multi-phosphoserine region of PER2 is therefore called the FASP domain. Notably, CK1ε^tau^ with a missense mutation that alters substrate recognition markedly decreases phosphorylation of the FASP domain, but increases phosphorylation of critical degradation sites (7, 16, 21).

PER proteins have two known CK1-regulated phosphodegrons. These phosphodegrons do not conform to standard CK1 recognition motifs but rather were identified by structure-function analysis (11, 22). CK1δ/ε phosphorylates a degron in PER1 at S122/S126 and also a degron in PER2 at S478/S482 (here called degron 1 and 2 or D1 and D2, respectively), to produce β-TrCP recognition motifs (11, 22, 23). β-TrCP binding leads to polyubiquitylation and proteasomal degradation of PER. Remarkably, the CK1ε^tau^ mutation that causes an 8-fold reduction of kinase activity on primed PER2 FASP increased phosphorylation of the PER2 degron 2 (7, 16, 21, 24). This ability of CK1 to phosphorylate both the FASP as well as the phosphodegrons, resulting in either stability or degradation of PER2, led to identification of the circadian phosphoswitch.

In the phosphoswitch, phosphorylation of the FASP domain stabilizes PER2 by inhibiting the ability of CK1 to phosphorylate degron 2 (21). This inhibition is due to direct binding of the phosphorylated FASP domain to conserved anion binding pockets in the CK1 substrate recognition domain, thereby blocking activity on degron 2 (20). While mutation of D1 has been reported to stabilize PER2 (23), the effect of the phosphoswitch on D1 has not yet been carefully examined. Because mice with homozygous mutation of the PER2 D2 degron had only a one hour lengthening of circadian period (25), the D1 degron may be an additional mechanism to regulate PER2 stability.

Here, building on advances in our understanding of the phosphoswitch, we studied the relationship between the two degrons in PER2. We asked if D1 activity is dependent on CK1, how D1 contributes to PER2 stability, and how two CK1-dependent degrons affect the phosphoswitch. Previous studies have established the utility of PER2 luciferase fusions in assessing protein stability with the help of real-time luminometry (25, 26). We made use of this system in combination with various PER mutants to investigate PER2 degradation independent of the canonical phosphodegron (D2) and CK1-binding. This alternate degron (D1) is also phosphorylated by CK1. We find that D1 motif in PER2 drives PER2 degradation but only when the canonical degron is mutated. While we found that the loss of CK1 binding decreased PER stability, PER2/PER1 homo and heterodimers allow bound CK1 to phosphorylate D2 in trans. This scaffolded phosphorylation of PER2 may explain the difference in degron utilization.

## Results

### A second CK1-regulated degron contributes to the PER2 phosphoswitch

To better understand the relative contributions of D1 and D2 in the phosphoswitch, we generated a series of mutants that inactivated the two degrons and FASP in a mouse PER2::Luc construct (Fig. 1A, 1B). Site-directed mutagenesis (SDM) was used to make serine-to-alanine substitutions at residues S93 in D1, S478 in D2 and S659 in FASP, both individually and in varying combinations (Fig. 1B). The half-lives of the various PER2::Luc mutants were then assessed in HEK293 cells as previously described (19, 21, 27). Because CK1 binds stoichiometrically to PER1 and PER2, in all experiments, either CK1ε or CK1ε^tau^ were co-expressed to ensure sufficient kinase for phosphorylation of the exogenous PERs.

**Fig. 1.**
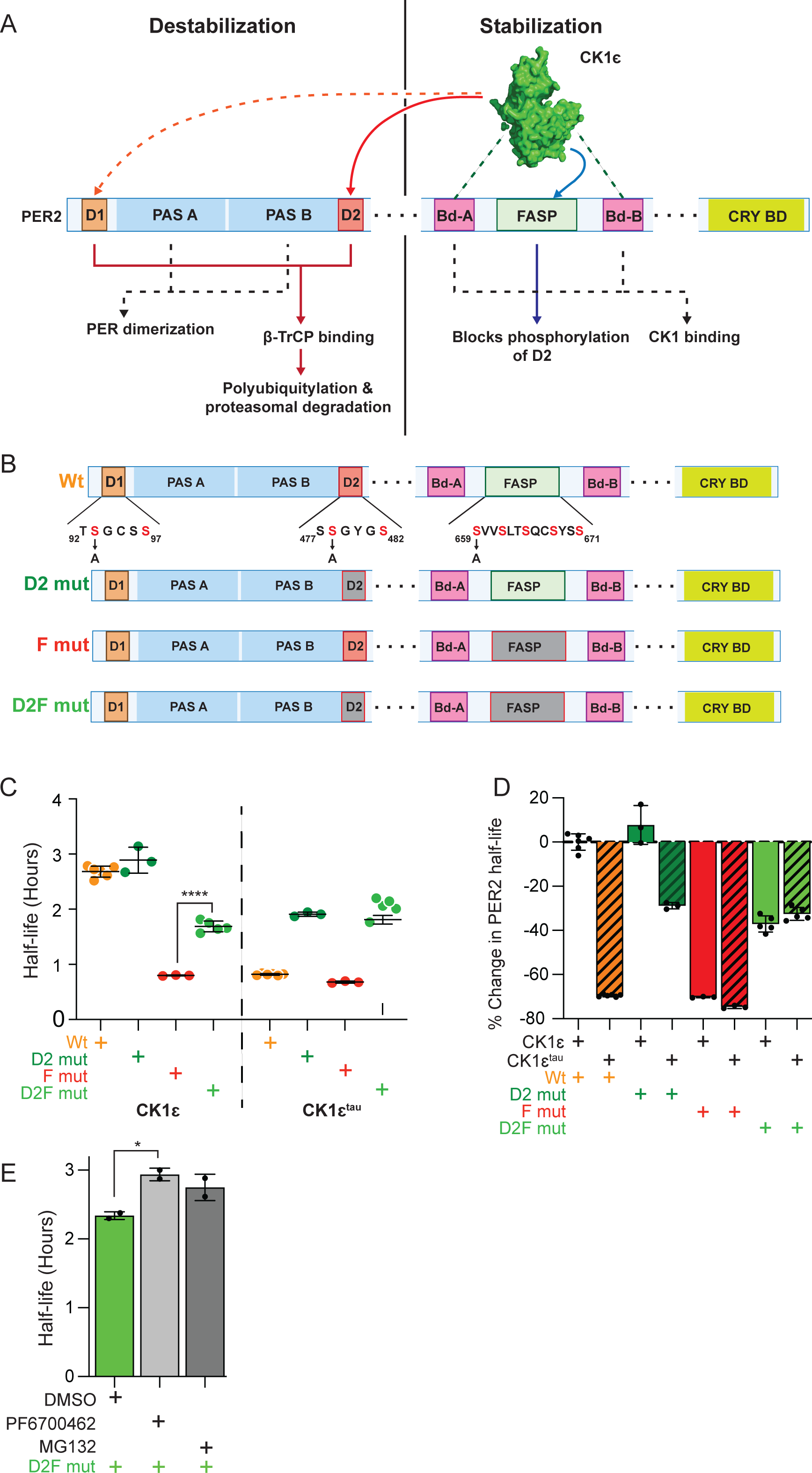
Additional CK1 regulated degron of the PER2 phosphoswitch. A) CK1 regulates the PER2 phosphoswitch. Binding to PER2 via CK1 binding domains is required for FASP phosphorylation which blocks degron 2 activation, resulting in PER2 stabilization (blue). CK1 phosphorylation of degron 2 allows for β-TrCp recognition and proteasomal degradation (red). Degron 1 is also recognized by β-TrCp but its phosphorylation by CK1 is undetermined (dotted orange). PAS domains A and B are required for PER dimerization as part of repressor complex formation. B) Mutation of PER2 phosphoswitch domains by alanine substitutions of phosphorylation-target serine residues S659A for the FASP, S93A for D1 and S478A for D2. C) CK1 continues to drive PER2 degradation despite loss of D2. Corresponding PER2 constructs were transiently expressed as shown, with CK1є or CK1є^tau^ (20 ng PER2 and 100 ng CK1є or CK1є^tau^ plasmids per 25 mm dish of HEK293). Protein translation was inhibited with 40 µg/mL cycloheximide. Points represent individual half-lives, with error bars indicating mean ±SD. Statistical significance was determined with one-way ANOVA. D) Overlapping phenotype of FASP mut and CK1є^tau^ is only partially rescued by mutation of Degron 2. Bars represent average percentage change in half-lives relative to Wt PER2 with CK1є. Error bars indicate ±SD. E) Activation of the putative non-D2 degron is sensitive to CK1 and proteasome inhibition. HEK293 transiently expressing D2F mut were incubated with 1 µM PF6700462 or 10 µM MG132 before translation inhibition. Bars represent average half-lives with error bars indicating ±SD. Statistical significance was determined by one-way ANOVA.

At baseline, CK1ε’s preference for phosphorylating the FASP domain limits the rate of PER2 degradation (24), and mutating Degron 2 (D2 mut) has an insignificant effect on PER2 half-life compared to Wildtype (Wt), ∼ 3 hours (Fig. 1C, 1D). As predicted by the phosphoswitch model, PER2 half-life is reduced 70%, to ∼ 1 hour when the FASP (F mut) is mutated, since CK1ε is then uninhibited and fully active for Degron phosphorylation. Similar to the FASP mutant, a 70% shorter, ∼1 hr half-life is obtained when CK1ε^tau^, which is inactive on the FASP, is expressed with Wt PER2. These results allowed us to test the relative importance of the D2 degron when CK1 is not inhibited by FASP phosphorylation. Notably, the Degron 2/FASP double mutant (D2F mut) showed only a 37% reduction in half-life, to around 2 hours (Fig. 1D) when expressed with CK1ε. In the presence of CK1ε^tau^, the D2 mutant similarly lengthened the half-life, but not to the level of wildtype PER2-luc. From this we conclude that Degron 2 is an important but not the only CK1-regulated degron in PER2. This is consistent with previous studies that demonstrated that mutation of D2 does not completely block PER2:β-TrCP interaction and only partially lengthens period in the mouse (21, 25).

To further test if this additional D2-independent mechanism was also CK1-dependent, we assessed the effects of CK1δ/ε inhibition on D2F mut stability. Treatment of D2F mut with the CK1δ/ε dual-inhibitor (PF6700462) at 1 µM lengthened its half-life by 0.6 hours. The 26S proteasome inhibitor (MG132) similarly slowed the degradation of D2F mut by 0.6 hours (Fig.1E). Thus, we inferred that an additional degron also uses CK1- dependent recruitment of the ubiquitin protein ligase complex.

A β-TrCP dependent degron N-terminal to the PAS domains was first identified in PER1 (Fig. 1A) (22, 28). This degron, now called D1, is conserved in PER2. D1’s contribution to PER2 stability has been assessed previously. Ohsaki concluded that D2 was the major determinant of PER2 stability. Reischl et al. similarly confirmed a role for D2 but with some data suggesting a role for D1 (23, 29). We elected to revisit the regulation using additional knowledge about the phosphoswitch. Thus, site-directed mutagenesis was used to generate mutants of D1 in various PER2 backgrounds (Fig. 1B).

### CK1ε preferentially drives PER2 degradation through D2 not D1

By comparing the percentage change in luciferase activity over time, we observed that the mutation of D1 alone did not produce any increase in PER2 stability when expressed with either CK1ε or with CK1ε^tau^ (Fig. 2A-C). However, when D2 was mutated to slow PER2 degradation, now the additional mutation of D1 slowed degradation further. Thus, D1 is functional as a degron only when D2 is inactive.

**Fig. 2.**
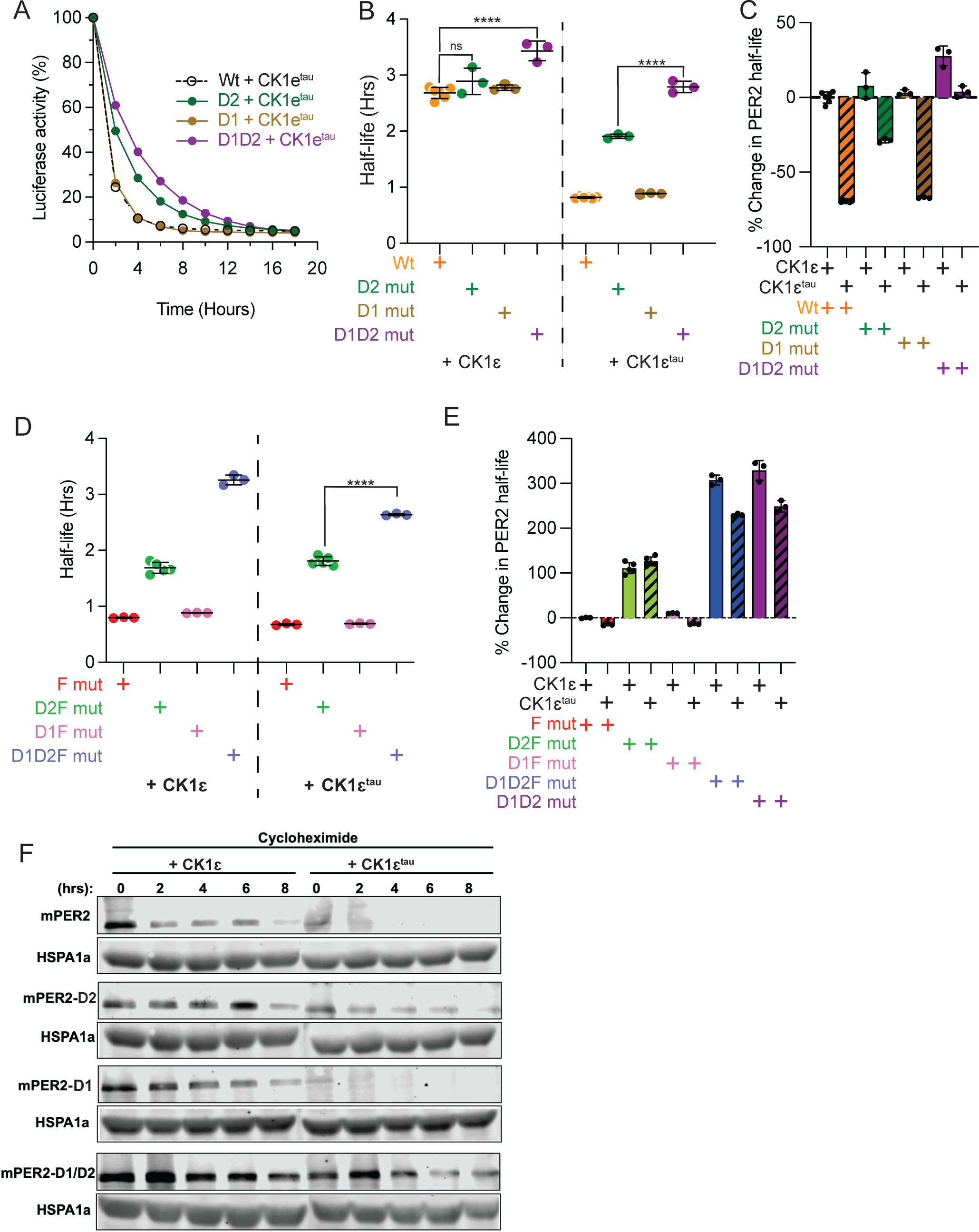
The second degron of PER2 is a back-up for the canonical degron, D2. A) Both WT (dashed) and D1 (brown) mutants are equally susceptible to CK1-driven degradation. Corresponding PER2 constructs were transiently expressed as shown, with CK1є or CK1є^tau^ (20 ng PER2 and 100 ng CK1є or CK1є^tau^ plasmids per 25mm dish of HEK293). Protein translation was inhibited with 40 µg/mL cycloheximide. Points represent average percentage of luciferase activity remaining with respect to 0 hours. ±SD was calculated and was too narrow to visualize. B) Simultaneous mutation of D1 and D2 offsets CK1є^tau^ enhanced degradation. Points represent individual half-lives, with error bars indicating mean ±SD. Statistical significance was determined with one-way ANOVA. C) D1D2 mut are significantly more stable than D2 mut. Bars represent average percentage change in half-lives relative to Wt PER2 with CK1є. D) Simultaneous mutation of D1 and D2 offsets F mut induced instability. Results are presented as in 2B. E) CK1є^tau^ enhanced degradation while reduced, persists with D1D2 mut. Bars represent average percentage change in half-lives relative to F mut PER2 with CK1є. F) Immunoblot of non-luciferase fused PER2’s recapitulated results of BDA. PER2 and CK1є or CK1є^tau^ were transiently co-expressed as indicated prior to Chx addition. Samples were harvested at 0, 2-, 4-, 6- and 8-hour timepoints before PER2 abundance was assessed with SDS-PAGE and Immunoblot.

Confirming the relative roles of D1 and D2, mutation of D1 (D1F mut) alone in the FASP mutant background failed to stabilize the protein, while double mutation of D1 and D2 extended the half-life ∼3-fold (Fig. 2D & E). In contrast, D2F mut had only a 1.3-fold increase in half-life. Thus, under the conditions of our assay, D1 appears to be masked by the greater activity of D2.

To test if this role of D1 was an artifact of the luciferase fusion construct, we measured the half-life of PER2 lacking luciferase by immunoblot over an 8-hour period after cycloheximide block of new protein synthesis (Fig. 2F). The loss of D1 function alone did not have a noticeable impact on PER2 abundance but the D1D2 mutant (mPER2-D1/D2) retained much higher protein levels than D2 mutant (mPER2-D2) alone.

These findings are consistent with the PER-luc half-life measurements and the importance of the D1 degron in PER2.

These data indicate that PER2 has at least two CK1-dependent phosphoswitch regulated degrons. Notably, the half-life of PER2 with mutation of both degrons D1 and D2 was still shortened by 30 to 45 minutes by CK1ε^tau^ expression (Fig. 2B-E). This suggested two possibilities. First, there might be additional CK1-dependent sites that also regulate PER2 stability. A second possibility builds on the known ability of CK1 bound to PER to phosphorylate PER binding partners such as CRY and CLOCK (30, 31). Thus, we tested if the dimerization of PER proteins allowed endogenous PER to provide a scaffold for CK1 to trans-phosphorylate degrons in the PER-luc constructs.

### Scaffolded phosphorylation of Degron 2

Models of CK1 regulation of the phosphoswitch have usually considered the interactions within each PER2:CK1 complex. However, PER proteins dimerize via their PAS domains, and a critical step of circadian clock regulation is the formation of large complexes containing dimerized PERs and bound CRYs (31, 32). The presence of a wildtype endogenous PER in the repressor complexes might explain how CK1ε^tau^ enhanced the degradation of PER with two mutant degrons. We hypothesized that endogenous PERs that heterodimerized with the ectopically expressed D1D2 mut are also phosphorylated by ectopic CK1ε^tau^, thus allowing binding of β-TrCP and subsequent polyubiquitylation of additional components of the repressor complex including D1D2 mutant PER2, accelerating their degradation. To test this model, we asked if a short half-life PER2 could shorten the half-life of a stable PER2. To do this, we co-expressed a FASP mutant PER2 lacking luciferase with a D1D2 mutant PER2-luc (Fig. 3A). Indeed, the PER2-FASP mutant (F mut (no luc)) co-expressed with D1D2 mut reduced its half-life by 0.7 hours (Fig. 3A), while a stable PER2 without luc (D1D2 mut (no luc)) had no effect. Similar half-life shortening was achieved when unstable PER2 (without luc) was co-expressed with D1D2F mutant PER2-luc. This suggested that PER2 stability is influenced by its dimerization partner.

**Fig. 3.**
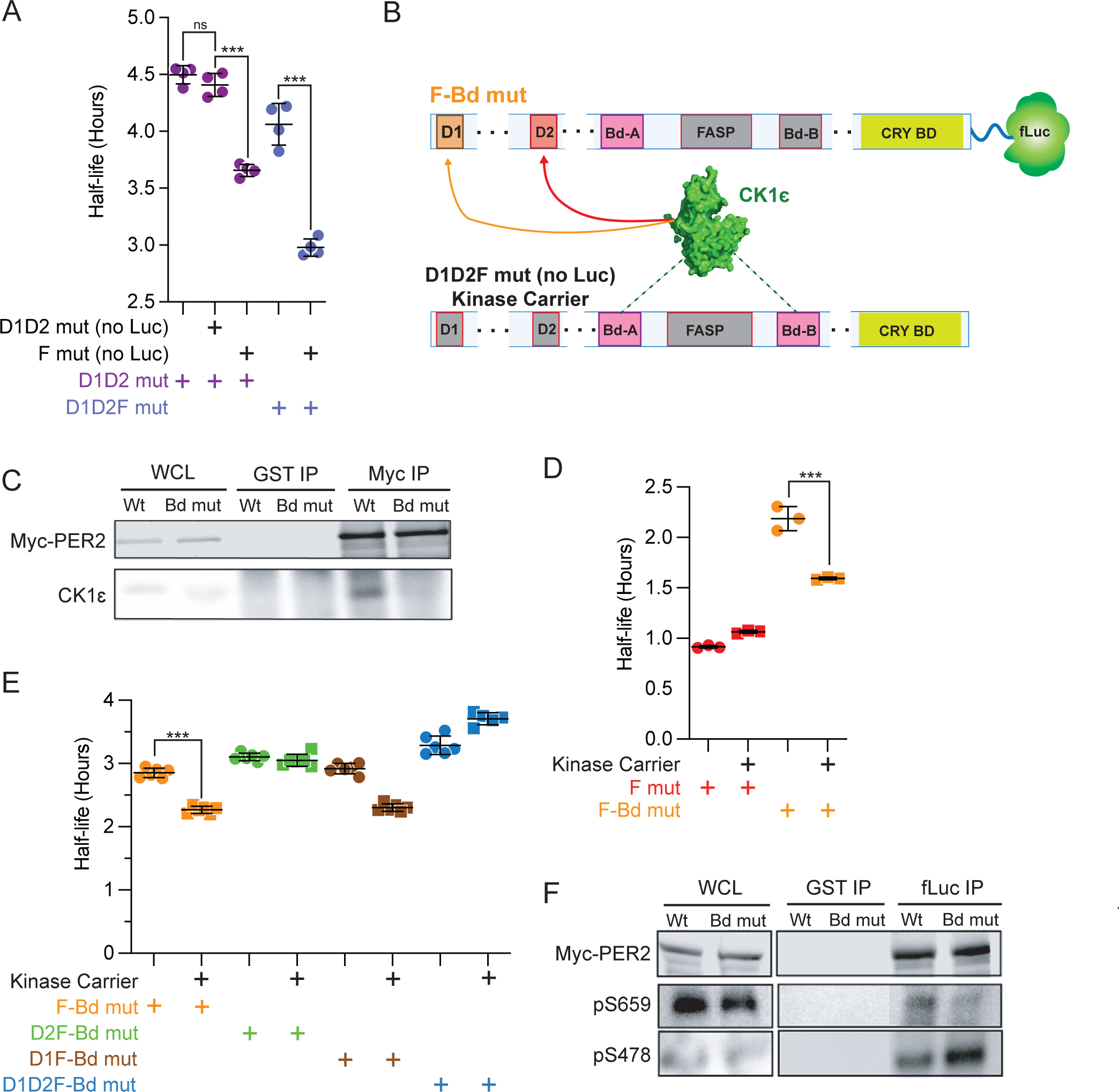
Phosphorylation of D2 facilitates PER2 degradation without direct binding of CK1Scaffolded. A) PER2 stability is influenced by its dimerization partner. Corresponding PER2 constructs were transiently expressed as shown, with CK1є (20 ng PER2::Luc, 20 ng PER2 non-luc and 100 ng CK1є plasmids per 25 mm dish of HEK293). For D1D2 mut and D1D2F mut only, 40 ng of Luc-fusion plasmids were transfected to control for total amount of PER2 plasmids. Protein translation was inhibited with 40 µg/mL cycloheximide. Points represent individual half-lives, with error bars indicating mean ±SD. Statistical significance was determined with one-way ANOVA. B) Cartoon indicates how CK1-binding domain mutants and Kinase carriers were used to test scaffolded phosphorylation. Boxes in gray indicate mutated domains. Kinase carriers have all phosphoswitch domains mutated and are not fused to luciferase. C) Bd-B mutation eliminated CK1-binding in PER2. Wt or Bd mut PER2 was expressed in HEK293. CK1-binding ability was assessed by immunoprecipitation of PER2 before SDS-PAGE and immunoblot for CK1є. WCL = Whole Cell Lysate. D) Scaffolded phosphorylation contributes to PER2 degradation. Corresponding PER2 and CK1є constructs were co-expressed with or without Kinase carrier (20 ng PER2::Luc, 20 ng Kinase carrier and 100 ng CK1є plasmids per 25 mm dish of HEK293). E) Scaffolded CK1 specifically targets D2. Experiment was performed as above. Points represent individual half-lives, with error bars indicating mean ±SD. Statistical significance was determined with one-way ANOVA. F) Loss of CK1 binding skews the phosphoswitch towards D2 activation. Wt or Bd mut PER2 was enriched via fLuc immunoprecipitation. After SDS-PAGE, immunoblotting was performed with antibodies raised against pS478 and pS659 peptides.

We hypothesized that the stable, degron-mutant PER2s (D1D2 mut and D1D2F mut) acted as a scaffold for bound CK1 to phosphorylate the degrons in its dimerization partner. To test if scaffolded phosphorylation allowed CK1 to phosphorylate a PER2 in trans in a PER2:PER2 dimer, we mutated the CK1 Binding-domain B (Bd-B) of PER2 to prevent CK1 binding to the PER2-luc fusion constructs (Fig. 3B), as recently described by An and colleagues (17). Co-immunoprecipitation analysis confirmed this mutation effectively eliminated the association of CK1ε with PER2 (Fig. 3C). The FASP Bd double mutants (F-Bd mut) were more stable than FASP mutants (half-life >2 hr versus <1 hr), likely because the homodimerization of the F-Bd PER2 results in repressor complexes that cannot bind CK1ε, thus preventing degron phosphorylation (Fig. 3D). We next generated a form of PER2 we called Kinase Carriers: PER2 without Luc and where D1, D2 and FASP sites were all mutant (D1D2F). In these stable Kinase Carriers, the phosphoswitch and degrons are inactivated and do not contribute to the regulation of repressor complex stability. As they have an intact PAS domain, they can still dimerize with PER2-luc and act as a scaffold for CK1 to phosphorylate other proteins, including any PER2 Bd mut in the repressor complex. Co-expression of F-Bd with Kinase Carrier partially restored its short half-life, halving the effect the Bd-B mutation had on F mut stability (Fig. 3D). Thus, we conclude that CK1 can regulate PER2 stability by phosphorylating both cis-bound PER2 and dimerized (trans) PER2.

CK1’s two modes of phosphorylating the PER2 degrons (in cis and in trans) may explain the difference between Degron 2 and Degron 1 utilization. To explore this, we assessed the stability of Degron 1 or Degron 2 mutants in a F-Bd mut background when co-expressed with Kinase Carrier and CK1ε (Fig. 3E). We saw that scaffolded phosphorylation promoted the degradation of F-Bd mut and D1F-Bd mut equally. This effect was lost when D2 was mutated (D2F-Bd mut), leading us to conclude that the primary target of scaffolded phosphorylation was D2, not D1. Counter-intuitively, in this same experiment D1D2F-Bd mut half-life increased slightly when stable Kinase Carrier was introduced, likely because this competed with dimerization with endogenous PER Finally, in comparison to Wt, Bd mutant PER2 had significantly reduced FASP phosphorylation (pS659) while D2 phosphorylation (pS478) was elevated when assessed by immunoblot (Fig. 3F). These findings agree with recently published studies that showed disruption of either Bd-A or Bd-B was sufficient to prevent phosphorylation of the FASP (17, 33).

### The phosphoswitch in PER1

Since repressor complexes incorporate varying combinations of PERs, we next assessed the role of the phosphoswitch as well as scaffolded phosphorylation on PER1 stability. Unlike PER2, PER1 does not possess Degron 2 (Fig. 4A). PER1 should thus behave similarly to PER2 D2 mut. By extension, we expected that mutation of the CK1 Bd would stabilize the protein, since Degron 1 is not a good target of scaffolded phosphorylation in PER2. As in PER2, mutating Bd-B of PER1 is sufficient to significantly reduce its association with CK1ε (Fig. 4D). As predicted, mutating its only degron (PER1-D1 mut) stabilized PER1. When the CK1 binding domain of PER1 (PER1-Bd mut) was mutated, the PER1 half-life also increased significantly but not as much as PER1-D1 mut (Fig. 4B). Next, we tested scaffolded phosphorylation of PER1 degron by co-expressing stable Kinase Carrier with PER1-Bd mut (Fig. 4C). The addition of Kinase Carrier increased the degradation of PER1-Bd mut, suggesting that unlike in PER2, Degron 1 was a target of scaffolded phosphorylation in PER1.

**Fig. 4.**
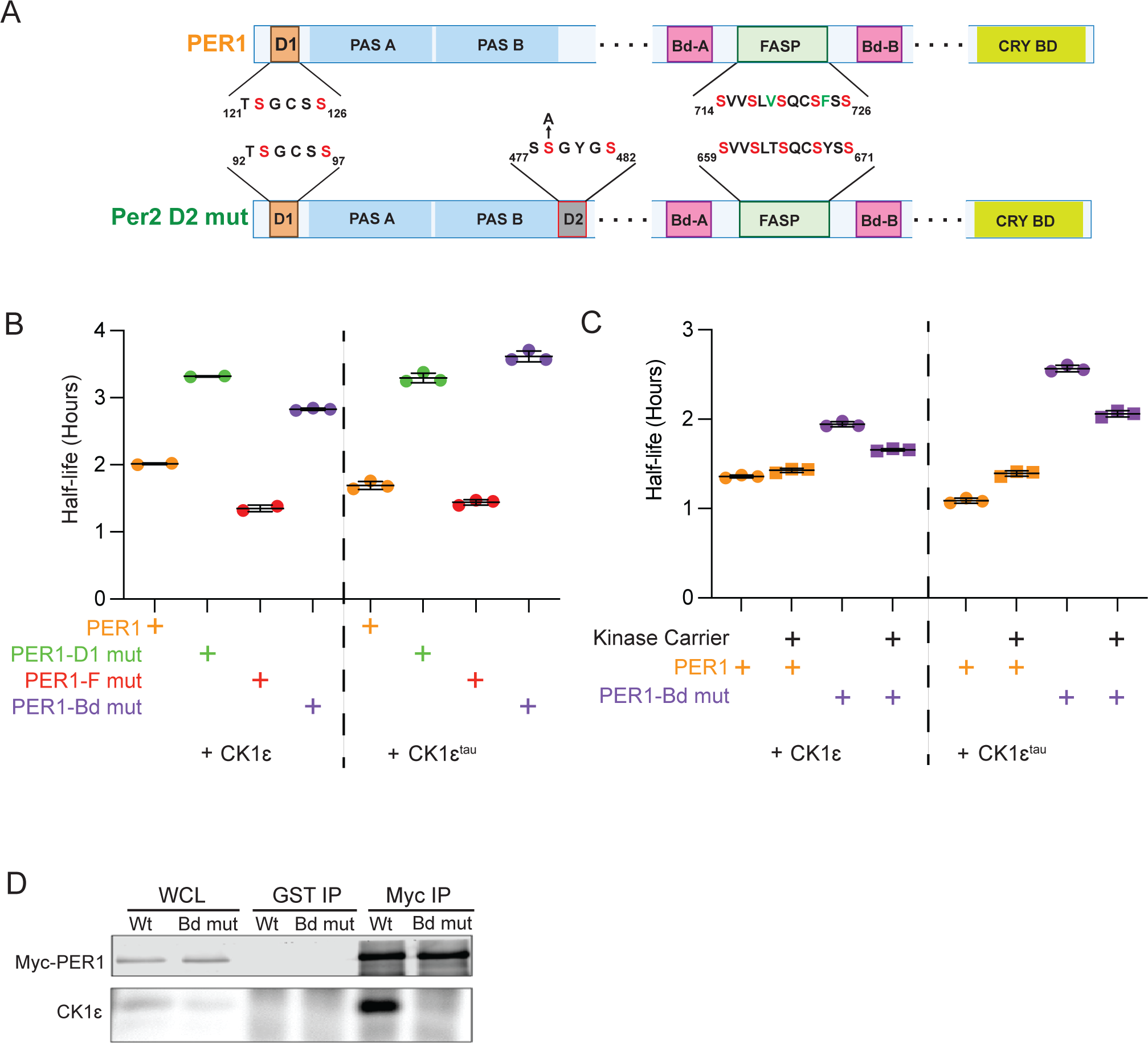
The phosphoswitch in PER1. A) Constructs for testing the PER1 phosphoswitch. B) Direct binding of CK1 is important for PER1 degradation. The halflife of the indicated PER1 constructs were determined as above. C) PER1 D1 has a limited capacity for scaffolded phosphorylation. The indicated PER1 constructs were co-expressed with or without Kinase carrier and in the presence of either CK1є or CK1є^tau^ (20 ng PER1::Luc, 20 ng Kinase carrier and 100 ng CK1є or CK1є^tau^ plasmids per 25 mm dish). Points represent individual half-lives, with error bars indicating mean ± SD. Statistical significance was determined with one-way ANOVA. D) Bd-B mutation eliminated CK1 binding to PER1. Wt or Bd mut PER1 was expressed in HEK293 and CK1 binding assessed by co-immunoprecipitation with PER1. WCL = whole cell lysate.

## Discussion

Our study identified additional levels of regulation of the PER2 phosphoswitch by CK1ε. While PER2 has two active degrons, Degron 1 was less efficient at driving PER2 degradation than the canonical phosphodegron D2 under the conditions of our assay. We confirmed that Degron 1 is regulated by CK1ε and the proteasome, similar to Degron 2. Mutation of both degrons was required to stop CK1-mediated decay. An and co-workers recently identified scaffolded phosphorylation in PER dimers (17). Here, we confirm and extend that finding to show that in PER homo- and heterodimers, CK1ε can phosphorylate PER2 Degron 2 and PER1 Degron 1 in trans.

Although the contribution of Degron 1 to PER2 stability has been previously suggested, its dependence on CK1 for phosphorylation and its regulation by the phosphoswitch has not been established (29). Mutations that prevent FASP phosphorylation generate strong effects on the circadian clock, shortening biological periods by as much as 4 hours. Degron 2 mutations on the other hand have comparatively weaker effects, extending biological periods by only about 1 hour. Since we have demonstrated a significant stabilization of PER2 in cells by mutating both degrons, we predict a double degron mutant would have a correspondingly longer biological period in vivo.

Amino acid sequence motifs that Casein Kinase 1 prefers to phosphorylate have been identified both in peptides and in intact proteins including PER, β-catenin, SV40 large T antigen, NFAT1, DARPP32, and APC (10, 11, 34–40). CK1 is most active on primed sites, where there is a phosphoserine in the -3 position (41). CK1 also phosphorylates biologically important unprimed motifs, but at significantly slower rates. For example, CK1 has much lower activity on the priming site S662 of hPER2, which lacks a well-characterized CK1 recognition sequence (24). Similarly, D1 and D2 are phosphorylated by CK1, but relatively slowly. Some measure of the relative activity of CK1 on the PER2 priming site versus D1 versus D2 can be estimated by the hierarchy of activity in phosphoswitch regulation. A second independent assessment of CK1 activity on these motifs can be made by looking at their substrate scores in a positional scanning peptide array (PSPA) analysis (Table 1) (39, 42). Both methods suggest the ranking of relative CK1 activity is PER2 primed>> PER2 priming>D2>D1. Thus, the sites that actually drive PER2 degradation are the most slowly phosphorylated sites.

**Table 1:**
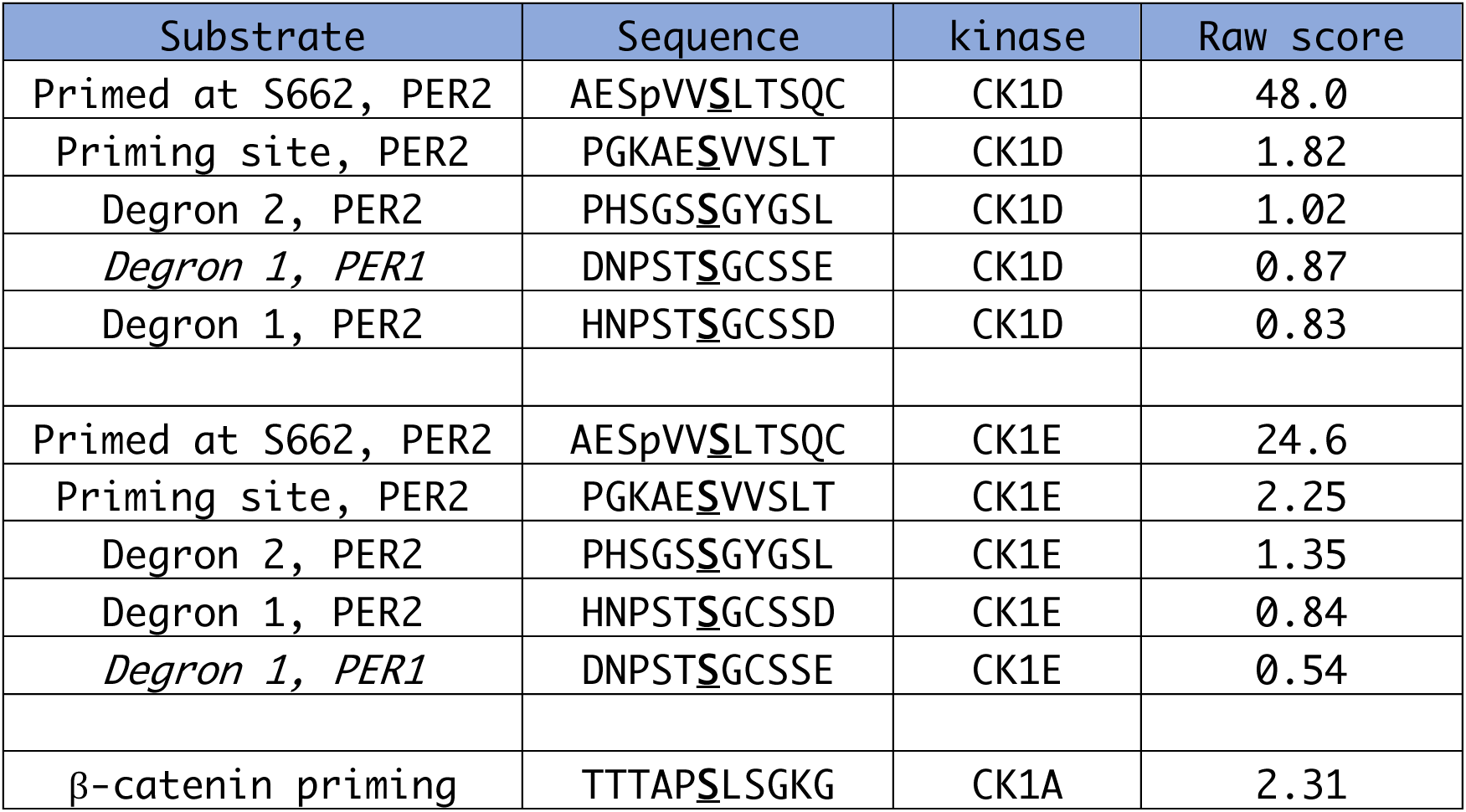
Scoring of substrate activity for CK1δ/ε with the Kinase library Site Score tool. β-catenin priming site score is included for comparison. Sp indicates phosphorylated site; bold underlined S indicates target residue. Data from Johnson et al., 2023 (39), using PhosphoSitePlus®(42).

Using peptide phosphorylation assays to predict kinase activity on proteins is complicated in the case of PER2 by its tight binding to CK1. Both degrons D1 (TSGCSS) and D2 (SSGYGS) resemble the canonical ß-TrCP motif DpSGɸXpS, and they score similarly in the PSPA analysis. In contrast, our experiments using full length proteins demonstrate that D2 has a much greater effect than D1 in regulating PER2 stability. The higher degradation activity of D2 that we observe may be due to greater accessibility of the bound kinase to the peptide in full length protein.

Our data as well as that of others suggest that PER2 is phosphorylated in the repressor complex both in cis and in trans through scaffolding of CK1 (17, 43). This scaffolded phosphorylation did not enable D1 activation in PER2, while it did allow D1 activation in PER1. This implies that steric considerations may allow CK1 greater access in trans to PER2 D2 than to D1 and that could contribute to the observed differences in activity between the degrons. This trans phosphorylation of D2 may drive PER degradation even when CK1 binding to one copy of PER is inhibited by various mechanisms, e.g., by competitive binding of CHRONO (43). Elucidating the mechanism of cis versus trans CK1 activity could be important in identifying new means of modulating circadian rhythms.

We developed a structural model to explain scaffolded PER phosphorylation. PER full length protein structures have not been solved, likely due to the presence of multiple intrinsically disordered regions. Predictive algorithms such as Alphafold do not fare much better outside of modelling the PAS domains and predicting several α-helices where CK1 and CRY binding domains are located. However, the structure of PER PAS domain homodimers is known (44, 45). In PER2, the 60 amino acid residues between Degron 1 and the start of the PAS domain are predicted to form α-helices (Fig. 5B). In PER1 this domain is predicted to be unstructured. Degron 2, on the immediate C-terminal side of PAS B, is immediately adjacent to the CK1 binding domains, and FASP site. We also know that CK1 must associate with both binding domains (pink) to access the FASP (Fig. 5C). Prior studies have shown that CK1 can phosphorylate D2 in the absence of the PAS domains (11). Dimerization also positions D2 of PER2 near the enzymatic cleft of CK1’ which is scaffolded on PER2’. On the other hand, D1 that is situated on a semi-flexible arm can bend back to access the natively bound CK1 but is not able to reach the scaffolded CK1’ (Fig. 5D). Applying the same techniques to PER1, we identified several features of the D1 arm in PER1, including high flexibility and a nine amino acid insertion immediately upstream of D1, which distinguish it from the D1 arm of PER2. Further study is required to determine if these differences are what enable scaffolded phosphorylation of Degron1 in PER1 but not PER2.

**Fig. 5.**
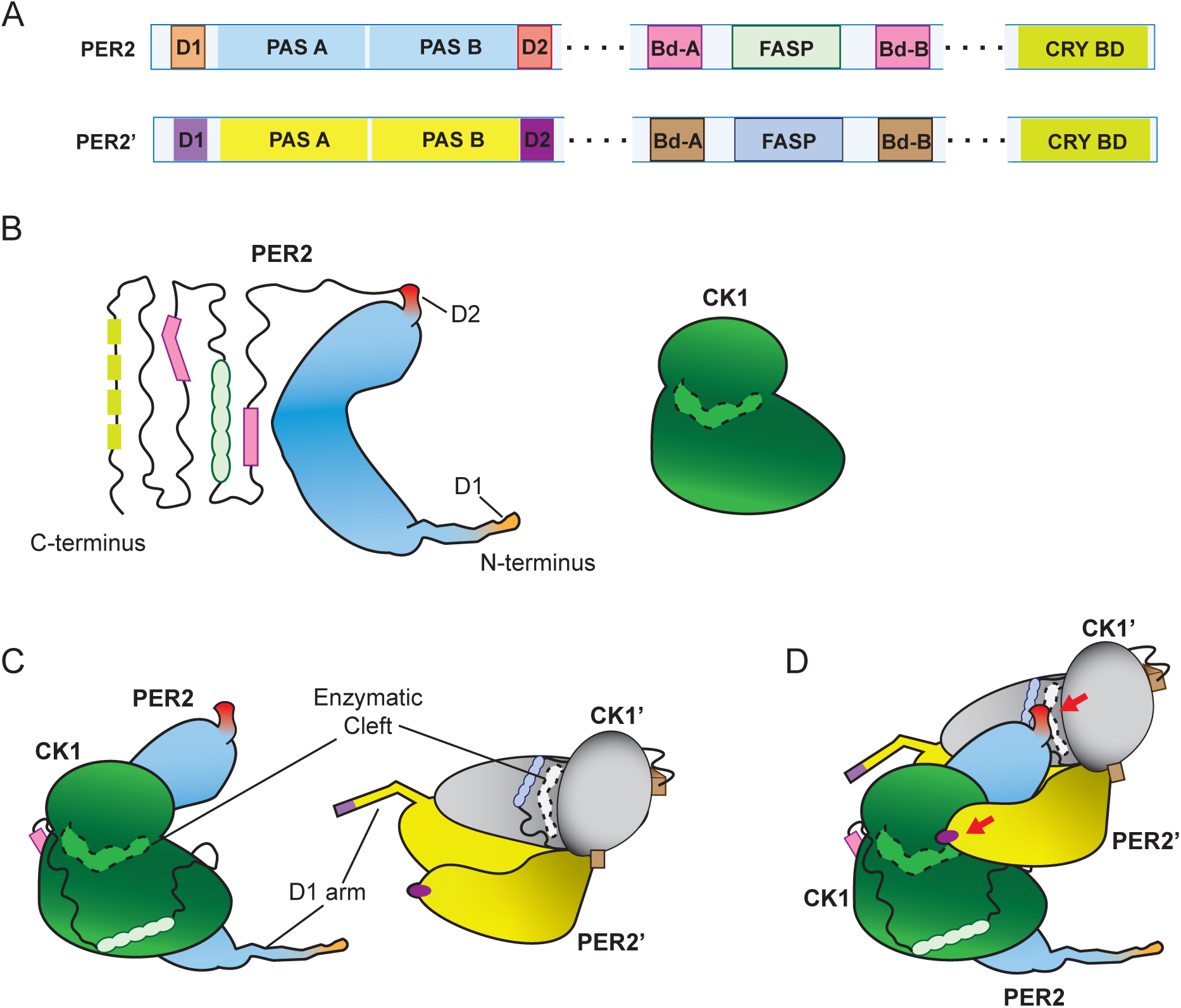
Predicted arrangement of unstructured PER2 domains inferred from PAS crystal structures and Alphafold models. A) Schematic of two identical molecules, PER2 and PER2’ that dimerize as part of a repressor complex. B) Cartoon of PER2 molecule and CK1. PAS domains (light blue) are highly organized with solved structures and are required for PER dimerization. D1 (orange) is situated on a semi-flexible arm N-terminal to the PAS domains. D2 (red) protrudes from the PAS domains and extends to the CK1Bd (pink) flanked FASP (light green), terminating in the Cry-binding domain. This extension is highly disordered and little structural information is currently available. Alphafold predicts alpha-helices at the D1 arm, CKBds and Cry-binding domain with confidence. C) Binding of PER2 to CK1 (green) and PER2’ to CK1 (grey). CKBd-A and B must both bind to CK1 forming a loop which exposes PER2 FASP to its enzymatic cleft (dashed outline). PER2’ and its bound CK1 are rotated to superimpose on dimerized PER2 PAS domains (3gdi). D) Dimerization of PER2 scaffolds their respective CK1s such that each PER D2 is accessible to the enzymatic cleft of its partner’s kinase.

In vivo, differential accessibility of the degrons to cellular phosphatases and deubiquitylating enzymes (DUBs) could further differentiate the roles played by each degron in regulating PER stability. We observed that D1 is positioned on a flexible arm extending N-terminally from the PAS domains of both PER1 and PER2 proteins. While the precise orientation of this arm relative to the full complement of proteins within the repressor complex remains unknown, it is plausible that the D1 region is more exposed to both phosphatases and DUBs. The infrequent phosphorylation of D1, coupled with its heightened vulnerability to dephosphorylation and deubiquitylation, may contribute to its lesser contribution to PER2 degradation.

We found that CK1-mediated decay of PER1 was congruent with the revised PER2 phosphoswitch model. Since PER1 only has the D1 degron, we expected PER1 to contribute less to repressor complex degradation than PER2. This is because in the PER2 phosphoswitch, D2 was demonstrated to have a dominant effect on stability over D1. Notably, Lee et al. observed that exogenous expression of PER2 generated more robust circadian rhythms than PER1 in PER deficient mouse endothelial fibroblasts due to lower troughs in PER2 protein abundance (2). Building on their finding that proper stoichiometry between positive and negative elements is essential for maintaining circadian rhythms, our study further suggests that the stoichiometry between PER1 and PER2 can fine tune its speed.

## Experimental Procedures

### Plasmid generation

mPER2 coding sequence was cloned into a pCS2-6xMycTag vector and firefly luciferase sequence was fused in frame to the C-terminus. FASP, Degron 1, Degron 2 and CK1ε binding domain mutants were generated via site-directed mutagenesis with KODone PCR Master mix (Toyobo). Constructs were amplified in DH5α E. coli and validated with Sanger sequencing.

### Cell culture and transfection

HEK293 cells were maintained at 37°C, 5% CO2 with Dulbecco’s Modified Eagle’s Medium (DMEM, Nacalai Tesque), 10% Fetal Bovine Serum (Hyclone) and 1% penicillin/streptomycin mixture (Gibco). 700,000 cells were seeded into a 35 mm dish (Corning) one day prior to transfection. The cells were transfected with 20 ng of PER and 100 ng of CK1ε expression constructs with pCS2 empty vector to a total of 1 µg plasmid with Lipofectamine 2000 (Invitrogen) as per manufacturer’s direction.

### Bioluminescence degradation assay (BDA)

Lumicycle media was prepared with phenol red-free high glucose DMEM (Life Technologies), 10% FBS and 1% penicillin/streptomycin. Luciferin was added just before use to a final concentration of 100 µM (Perkin Elmer). 20 hours after transfection, the cells were incubated with 40 µg/mL cycloheximide (MPBio) to inhibit protein translation. Luciferase activity was recorded in real-time with the Lumicycle (Actimetrics). The half-life of PER constructs was determined with non-linear regression analysis using GraphPad Prism software.

### Assessing PER degradation via immunoblot

Transfection and cycloheximide addition were performed as in BDA protocol. Samples were harvested at 0, 2-, 4-, 6- and 8-hour timepoints with cell lysis buffer (1% Nonidet P-40, 0.5% sodium deoxycholate, 50 mM Tris hydrochloride, 150 mM sodium chloride in water). PhosStop (Roche) and Complete protease inhibitors (Roche) were added to the lysis buffer just prior to use as per manufacturers direction. Cell lysates were denatured by boiling with 3X SDS Blue Loading Buffer (New England Biolabs). Samples were resolved on SDS-PAGE gel and transferred onto nitrocellulose membranes (Millipore, Immobilon). The mouse monoclonal anti-Myc Tag antibody, clone 4A6 (Merck Millipore, #05-724) was used to detect 6Mt-mPER2 while rabbit polyclonal antibodies against HSPA1a (Invitrogen, PA5-34772) was used as a loading control. Dylight 680 nm conjugated goat anti-rabbit or -mouse IgG (Invitrogen) were used as secondary antibodies. Immunoblots were imaged with Odyssey Imager (Li-Cor).

### Immunoprecipitation of PERs

Cells were transfected and harvested as described above. 75 µg of protein lysate was used for each reaction. Pulldown of myc-PER1 or myc-PER2 constructs was accomplished with Protein G Dynabeads (Invitrogen) and 2 µg of anti-c-myc (4A6; Merck Millipore). For immunoprecipitation of PER2::Luc constructs rabbit anti-firefly luciferase IgG (Abcam, ab21176) was used instead. Pulldown with anti-GST antibody (z-5, Santa Cruz) was performed as a control. The lysates and antibody:Dynabead mixtures were incubated at room temperature for 30 minutes with rotation. Dynabeads were separated from the supernatant with a magnetic stand and the supernatant was discarded. The beads were then washed thrice with PBS-T (0.1% Tween-20). They were then transferred to a clean tube for elution by boiling in 1X SDS Blue Loading Buffer (New England Biolabs) at 85°C for 10 minutes. Immunoblot of the samples was performed as described above. Polyclonal antibody against CK1ε (H-60, Santa Cruz) was used for these experiments.

### Immunoblotting of PER2 pS478 and pS659

Antibodies for detection of PER2 pS478 and pS659 were generated and used as described previously (19). Goat anti-rabbit HRP conjugate (Santa Cruz) was utilized as a secondary antibody and signal detected with ECL western blotting substrate (Pierce) on a ChemiDoc (Biorad).

## Acknowledgements

We thank members of the Virshup Lab, especially Rajesh Narasimamurthy and Ewan Stephenson for the stimulating discussions around the Clock and also Yunka Wong for his support in the lab. This work was funded by the National Medical Research Council of Singapore Grant MOH-000600 (to D.M.V.).

## Data Availability

All data are contained within the manuscript.

